# Immobilization, Direct Electrochemistry and Electrocatalysis of Hemoglobin on Peptide-Carbon Nanotube Modified Electrode

**DOI:** 10.1101/2020.06.16.153767

**Authors:** Mohammadali Sheikholeslam, Piyush Nanda, Mark Pritzker, P. Chen

**Affiliations:** Department of Chemical Engineering, University of Waterloo, 200 University Avenue West, Waterloo, Ontario N2L 3G1, Canada; Waterloo Institute for Nanotechnology, University of Waterloo, 200 University Avenue West, Waterloo, Ontario N2L 3G1, Canada; Biosensor Research Center, Department of Biomaterials, Tissue Engineering and Nanotechnology, School of Advanced Technologies in Medicine, Isfahan University of Medical Sciences, Isfahan 81746-73461, Iran

## Abstract

The direct electrochemistry of hemoglobin (Hb) immobilized on a glassy carbon electrode (GCE) by mixing with a hybrid dispersion of self-assembling peptide EFK8 and single-walled carbon nanotubes (SWNT) was investigated and shown to have effective properties as a biosensor for hydrogen peroxide (H_2_O_2_). Preliminary experiments involving cyclic voltammetry (CV) and electrochemical impedance spectroscopy (EIS) showed that the presence of SWNTs in the modifying peptide layer on GCE significantly enhanced the electrochemical response of the electrode for the ferricyanide/ferrocyanide redox couple. Also, when more layers of the peptide-SWNT dispersion were applied to the surface, the electrochemical response of the GCE was further increased. This behavior was then exploited to fabricate a biosensor for H_2_O_2_ by mixing hemoglobin with the hybrid peptide-SWNT dispersion and casting this mixture on a glassy carbon electrode. AFM imaging, CV and EIS analysis revealed successful immobilization of Hb on the electrode and the ability of the electrode to enable direct electron transfer from Hb to the electrode surface. In particular, CVs obtained in 0.1M PBS (pH 7.0) showed that the immobilized Hb retained its bio-catalytic activity for Fe ions, indicating that it had not denatured and the hybrid layer remained biocompatible with Hb. Furthermore, this electrode was found to accurately measure the H_2_O_2_ concentration over the range from 20 to 9.6×10^2^ μM in 0.1M PBS (pH 7.0) under mediatorless conditions using both CV and amperometry techniques. In this way, the EFK8-SWNT hybrid layer shows promise as a biocompatible layer for simple non-covalent enzyme immobilization and the basis of a mediatorless biosensor.

## 1. Introduction

Since the 1970s, direct electron transfer between redox proteins and electrode surfaces and its application for mediatorless electrochemical biosensors have been extensively studied. The most efficient method to make a redox protein-based electrochemical biosensor is to establish direct electron transfer between the protein and electrode. The use of a mediator facilitates not only the electron transfer between the electrode and enzyme but unfortunately also various interfering reactions. Mediatorless biosensors can offer better selectivity since they are able to operate in a potential range closer to the redox potential of the protein itself and thus make interfering reactions less likely to occur [1][2]. In addition, most *in-vivo* devices are mediatorless to prevent the potential of leaching and toxicity of the mediator. Mediated systems also tend to be less stable during extended continuous operation [3]. Normally direct adsorption of proteins on the electrode surface leads to their denaturation and loss of their catalytic and electrochemical activity [4]. Thus, an important first step in designing a mediatorless biosensor is to immobilize the protein without denaturation.

Hemoglobin (Hb) is an important protein in red blood cells as a reversible oxygen carrier in the body through its four polypeptide chains which contain electroactive heme groups. It has been reported that Hb also can catalyze the reduction of hydrogen peroxide [5]. The ability to rapidly and accurately determine hydrogen peroxide concentration is very important since it is the product of many enzymatic reactions and is also commonly found in food, clinical, pharmaceutical, industrial and environmental systems [5]–[7][2]. A number of techniques such as HPLC [7], titrimetry [8], spectrometry [9], chemiluminescence [10] and electrochemical methods [11] are currently available for hydrogen peroxide determination.

Electrochemical methods for hydrogen peroxide analysis have attracted extensive interest because they are fast, less prone to interferences and relatively inexpensive [7]. Basically, two types of amperometric enzyme-based H_2_O_2_ probes can be used – mediated biosensors and mediatorless biosensors. Although the mediated H_2_O_2_ biosensors can detect very low concentrations by use of electron transfer mediators such as ferrocene derivatives [12], hexacyanoferrates [13], tetrathiafulvalene [14] or phenazine methosulphate [15], the danger exists that mediator molecules can contaminate the sample or electrode system or can diffuse out of the enzyme layer [16]. Mediatorless biosensors which operate through the direct electron transfer between redox proteins and electrode have gained increasing attention because they do not suffer from this leakage problem and have potentially simpler design without the need for a chemical mediator.

In addition to its ability to electrocatalyze H_2_O_2_ reduction, Hb is an ideal model protein for the study of the direct electron transfer of heme molecules due to its commercial availability, reasonable cost and well-known structure. However, the direct electrochemical reaction of Hb on an electrode does not readily occur for a number of reasons. Since the heme group resides inside the Hb structure and is surrounded by the polypeptide chains, the electrons being transferred must travel a large distance to the electrode surface and a mediator is often required to help transport them. Once adsorbed, Hb on the surface frequently denatures and loses its electrochemical activity and bioactivity. Hb becomes a barrier to electron transfer on the electrode surface once it becomes denatured [17]. Finally, Hb located at the electrode surface may not have a favorable orientation for electron transfer. To date, different methods and nanomaterials including Au, Pt and CdTe nanoparticles as well as graphene, CNTs and TiO2 nanorods have been employed to overcome the above obstacles and facilitate direct electron transfer from Hb to the electrode [5], [7], [18]–[29]. These methods mainly operate by providing a strong electronically conducting environment for

Hb molecules on the surface. Although promising results have been observed using these nanomaterials, the biosensor efficiency of Hb–based H_2_O_2_ biosensors is still lower than other ones using different types of heme proteins such as horseradish peroxidase (HRP) and catalase (CAT) [2].

Recently, the potential of using ionic-complementary peptides in biocompatible electrodes to immobilize glucose oxidase covalently and fabricate glucose biosensors was demonstrated [30][31][32]. These peptides can self-assemble on surfaces as β-sheet layers that have considerable biological and physiological stability [33][34] and good *in vitro* and *in vivo* biocompatibility [35][36]. These are crucial properties for an *in vivo* biosensor. However, at the same time, these results showed that the peptides on the surface act as barriers to electron transfer from the enzyme to the electrode and thereby hinder the electrode response and biosensor sensitivity. Previously, we have shown the ability of the same self-assembling peptides to disperse multi-walled and single-walled carbon nanotubes in water [37], [38]. The surfaces modified with the peptide-CNT dispersions also have enough biocompatibility for cell attachment and growth [37]. Consequently, we propose that the combination of the peptides and carbon nanotubes in the form of hybrid coating on the surface should provide a biocompatible electrode for protein immobilization and *in vivo* biosensing with an enhanced electrochemical response. This combination can provide a basis for the design of a mediatorless biosensor.

In the current study, we report a novel yet simple method to modify and increase the electrochemical response of a glassy carbon electrode using a self-assembling peptide-SWNT hybrid coating. Then we show that this coating can adsorb Hb physically, remain biocompatible enough for Hb to retain its redox activity and thereby enable direct electron transfer from the Hb to the electrode and improve its catalytic activity toward H_2_O_2_ reduction.

## 2. Materials and Methods

### 2.1 Materials

The ionic complementary peptide EFK8 with a sequence of FEFEFKFK where F corresponds to phenylalanine, E to glutamic acid and K to lysine was used in this study. This peptide was synthesized in our laboratory using an Aapptec Apex 396 peptide synthesizer (Aapptec LLC, USA). At neutral pH, F is a neutral hydrophobic residue, while E and K are negatively and positively charged, respectively. The peptide was protected by acetyl and amino groups at the N terminus and C-terminus, respectively, to prevent end-to-end electrostatic interactions between peptides. The metallic SWNTs (carbon > 90%, carbon as SWNT > 77%) were purchased from Sigma-Aldrich Co (catalog# 727777, lot# MKBH7136V). Hemoglobin (from bovine blood), potassium ferrocyanide (K_3_Fe(CN)_6_) and H_2_O_2_ (30%) were obtained from Sigma-Aldrich. Phosphate buffer saline (PBS) solution was made using Na_2_HPO_4_ and NaH_2_PO_4_ both at analytical grade (EMD).

### 2.2 Methods

#### 2.2.1 Peptide synthesis

All amino acids (Fmoc protected), activator 2-(6-chloro-1H-benzotriazole-1-yl)-1,1,3,3-tetramethylaminium hexafluorophosphate (HCTU) and Rink Amide-AM resin were obtained from Aapptec LLC (USA). All other solutions were purchased from Acros Organics (USA). The EFK8 peptide with a molecular weight of 1162.60 g/mol was synthesized in our laboratory via the solid-phase peptide synthesis (SPPS) method using an Aapptec Apex 396 peptide synthesizer before being purified by repeated precipitation in cold ether. Then it was freeze-dried and the final powder stored in a refrigerator. Matrix-assisted laser desorption ionization time-of-flight mass spectroscopy (Q-TOF Ultima Global, Waters, Milford, MA, USA) was used to measure the molar mass of the synthesized peptide.

#### 2.2.2 SWNT dispersion preparation

The stock suspensions were prepared by combining EFK8 and as-received SWNT (carbon > 90%, carbon as SWNT > 77%) together at a 1:1 mass ratio in pure water (18.2 MΩ; Millipore Milli-Q system) to yield concentrations of 2 mg/ml of both the peptide and SWNTs. The concentration of 2 mg/ml SWNT is based on the mass of as-received material and so may include components that are not actual SWNT. The suspensions were then mixed for 1 hour using a Qsonica XL-2000 probe sonicator at a power of 10W and centrifuged at a speed of 2,000×g for 1h to produce a supernatant that was decanted for later use in electrode modification.

#### 2.2.3 Electrochemical analysis

Electrochemical measurements were done using a VMP3 potentiostat/ galvanostat (Bio-Logic instruments) and a three-electrode electrochemical cell at room temperature. Glassy carbon (GCE, diameter: 5 mm) was used as the working electrode, platinum wire as the counter electrode and saturated calomel electrode (SCE) as the reference electrode. Electrolytes were deoxygenated with pure nitrogen before each experiment. Cyclic voltammetry was conducted at a scan rate of 100 mVs^−1^ over the potential range from 0.6 V to −0.2 V in 10 mM K_3_Fe(CN)_6_ dissolved in 1M KCl electrolyte, from 0.2 V to −0.8 V in 0.1 M PBS and from 0 V to −0.8 V in 0.1M PBS (pH 7.0). Electrochemical impedance spectroscopy (EIS) was performed potentiostatically with 10 mV amplitude sinusoidal waves superimposed on a constant potential of 0.29 V over a frequency range of 1 mHz − 100 kHz in 10 mM K_3_Fe(CN)_6_ dissolved in 1M KCl electrolyte. Amperometric experiments were carried out in 0.1M PBS (pH 7.0) containing different concentrations of hydrogen peroxide.

#### 2.2.4 Scanning electron microscopy

The EFK8-SWNT coatings were carefully removed from the GCE surface in entirety with the use of a polishing cloth and examined with SEM. The SEM images were obtained using an FE-SEM (LEO 1530) unit.

#### 2.2.5 Electrode modification

Before modification, the electrode was polished successively with 1.0, 0.3 and 0.05 μm alumina slurry on a polishing cloth to obtain a mirror-like surface followed by bath sonication in ethanol and water. Then the electrode was washed with acetone and air-dried. Electrode modification was done by casting 35 μl of the peptide or hybrid dispersion on the GCE surface and then left to air-dry. For hemoglobin immobilization, Hb was dissolved in water and then mixed with an EFK8-SWNT dispersion at a 1:1 (v/v) ratio. Then the mixture was cast on the GCE surface as mentioned above and left overnight to dry at 4°C.

#### 2.2.6 Atomic force microscopy

AFM images in the PeakForce^®^ QNM mode were collected of GCE surfaces modified with EFK8-SWNT and EFK8-SWNT-Hb using a Dimension Icon^®^ AFM (Bruker Nano Surfaces, Santa Barbara, CA). The samples were prepared by incubating 10μl of EFK8-SWNT or EFK8-SWNT-Hb suspensions on a GCE overnight. Silicon nitride tips (type SCANASYST-AIR, Bruker) with a radius of 2 nm were used for AFM imaging.

## 3. Results and Discussion

### 3.1 Electrode modification using EFK8 and EFK8-SWNT hybrid dispersions

As the first step, we investigate the effects of the following two modifications of the GCE – first with a coating formed from an EFK8 peptide solution and the second with an EFK8-SWNT dispersion coating. Cyclic voltammetry was conducted on these modified electrodes in a 1M KCl solution containing 10 mM K_3_Fe(CN)_6_. The resulting current associated with the [Fe(CN)_6_]^3−^/[Fe(CN)_6_]^4−^ couple is used as a measure of the electrochemical response of the electrodes. A comparison of the scans obtained on the bare GCE and EFK8-modified GCE shows that the presence of the peptide alone on the surface diminishes the cathodic and anodic peak currents associated with the [Fe(CN)_6_]^3−^/[Fe(CN)_6_]^4−^ couple (Figure 1). This is presumably due to the peptide layer blocking the electron transfer pathway between the solution and the electrode. On the other hand, when the GCE is coated with EFK8-SWNT, the electrochemical response is improved and the current associated with the [Fe(CN)_6_]^3−^/[Fe(CN)_6_]^4−^ couple increases significantly. This increase can be attributed to the presence of highly conductive SWNTs on the surface that facilitate electron transfer to the electrode. The separation between the anodic and cathodic peaks (E_Pa_ – E_Pc_) obtained on the EFK8-SWNT-modified GCE is measured to be 59.2/n mV (where n = 1 in this case), which is reflective of a reversible reaction.

**Fig. 1.**
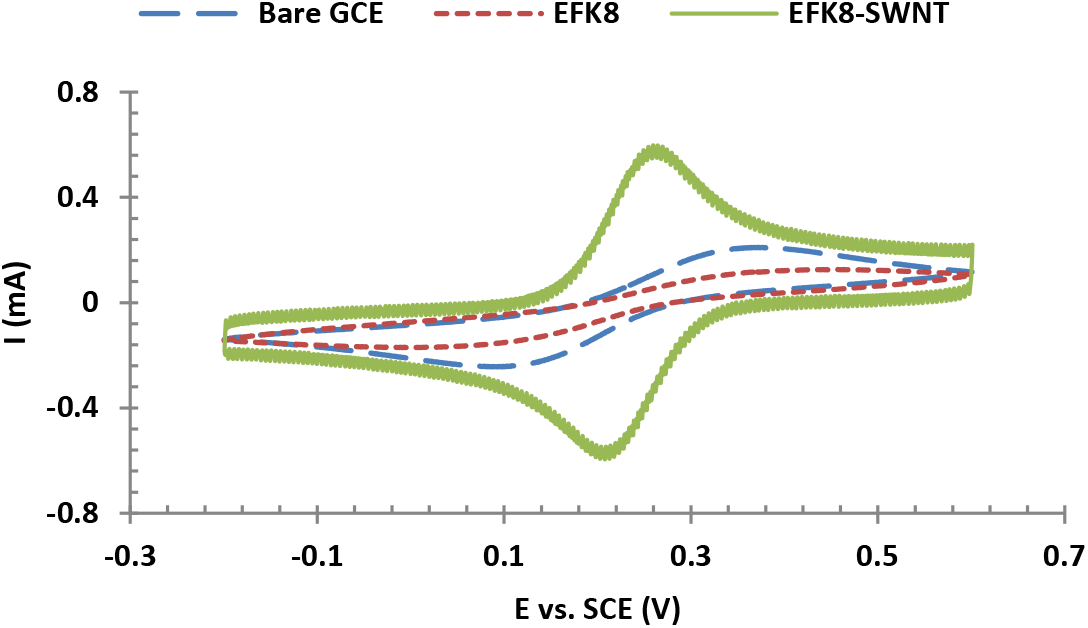
CVs obtained on bare GCE, GCE modified with EFK8 and GCE modified with EFK8-SWNT in 1M KCl containing 10 mM K_3_Fe(CN)_6_ at a scan rate of 100 mVs^−1^.

To further characterize the electrochemical response of the electrodes, electrochemical impedance spectroscopy was conducted. When the data are plotted in the form of a Nyquist plot, the diameter of the semicircle obtained at higher frequencies corresponds to the polarization resistance, which is a reflection of the kinetics of the electron transfer reaction at the electrode surface. A comparison of the responses in Figure 2 shows that the impedance of the EFK8 film is always larger than that of the EFK8-SWNT coating at the same frequency, consistent with expectations if EFK8 alone is a greater barrier for electron transfer at the electrode surface. Again, this observation is another indicator of the positive role of SWNTs in tunneling electrons to the surface.

**Fig. 2.**
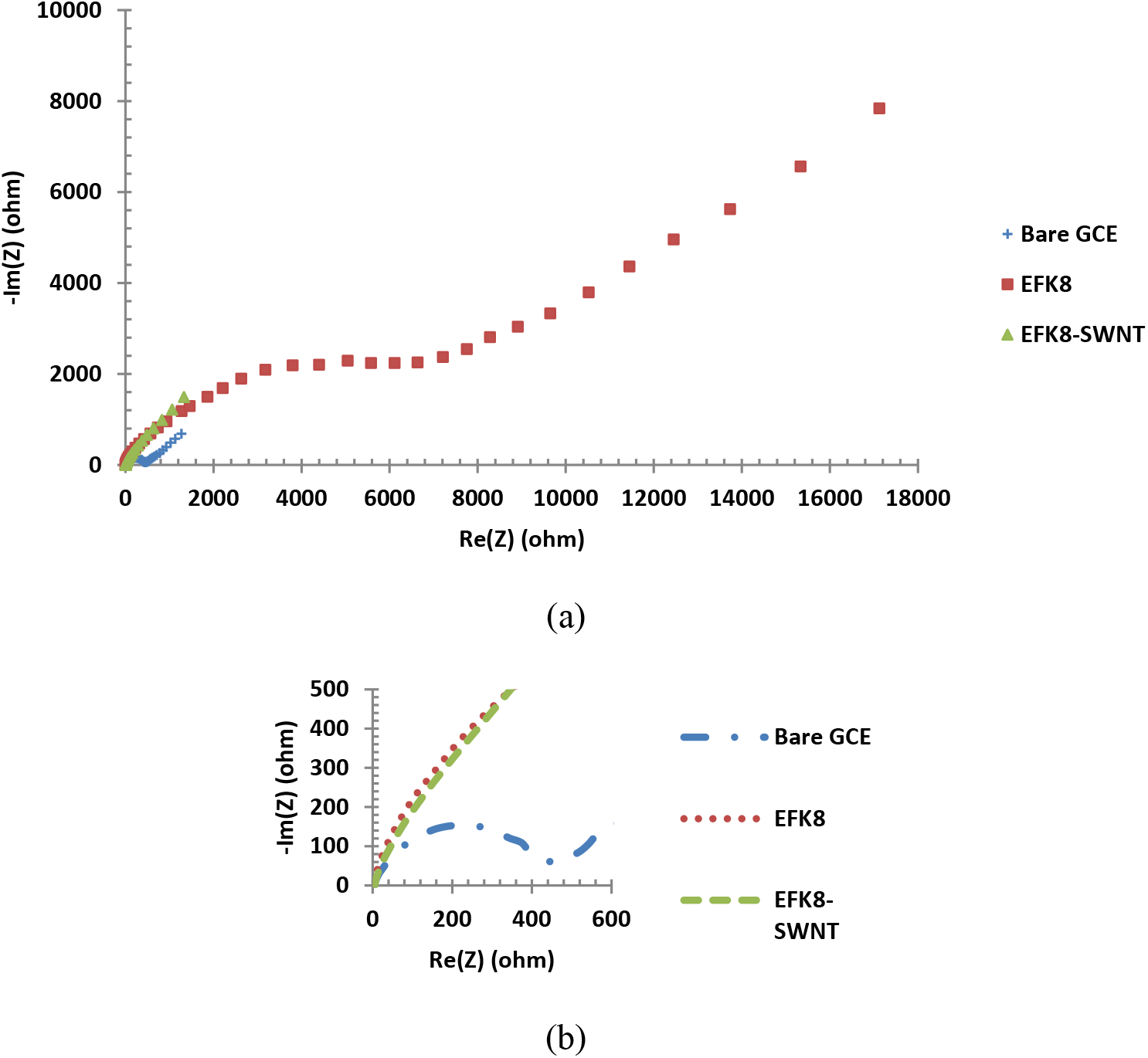
(a) Nyquist plots obtained on bare GCE, EFK8-modified GCE and EFK8-SWNT-modified GCE over a frequency range of 1 mHz – 100 kHz in 10 mM K_3_Fe(CN)_6_ dissolved in 1M KCl electrolyte. (b) Expansion of Nyquist plot in higher frequency region.

Cyclic voltammetry of the EFK8-SWNT-modified GCE has also been conducted over a range of scan rates between 25 and 200 mV s^−1^. Analysis of these CVs reveals that the peak currents for the [Fe(CN)_6_]^3−^/[Fe(CN)_6_]^4−^ couple rise linearly as the scan rate increases (Figure 3a), while the peak separation remains constant and the cathodic and anodic peak potentials change by less than 1%. Also, the peak currents and potentials do not change over 50 consecutive scan cycles, indicating that the hybrid coatings remain stable (Figure 3b).

**Fig. 3.**
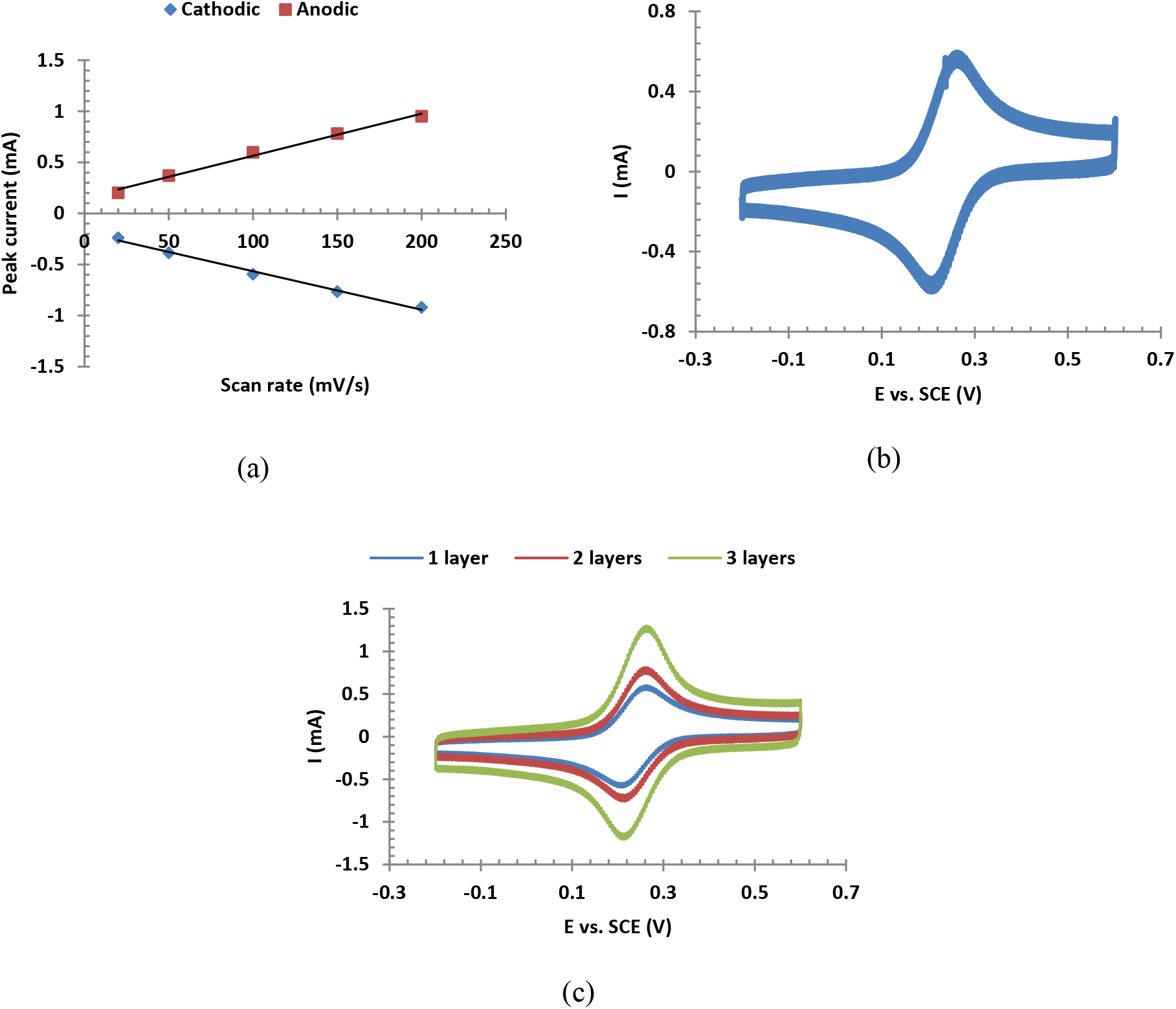
(a) Effect of scan rate on the cathodic and anodic peak currents of Fe(CN)_6_^3−^/Fe(CN)_6_^4−^ couple on an EFK8-SWNT modified electrode. (b) Traces of the CV responses over 50 cycles. (c) Effect of the number of EFK8-SWNT hybrid layers on the CVs obtained at a scan rate of 100 mV s^−1^ in 1M KCl containing 10 mM K_3_Fe(CN)_6_.

In order to investigate the effect of the thickness of EFK8-SWNT coatings, additional layers (each of which is introduced as a 35 μl dispersion) were applied to the GCE surface after drying the previous coating. Although the addition of more layers might be expected to impede electron transfer, the opposite is actually observed. As can be seen in Figure 3c, the presence of more layers increases the peak current probably due to an increase in surface area and the amount of SWNT on the surface. The SEM image in Figure 4a presents a top view of the EFK8-SWNT coating showing that it consists of fibers laid over each other. When this coating was removed from the GCE electrode, a crack in the coating developed. This turned out to be fortuitous since it reveals some of the EFK8-SWNT structure that would not otherwise be evident. As shown in Figure 4b and at higher magnification in Figure 4c, the EFK8-SWNT coating has a fibrous structure, consistent with that observed in Fig. 4a. The SWNTs cannot be distinguished from the peptide fibers due to similarities in their size and composition and the fact that the SWNTs are normally wrapped with peptides in the dispersion. Nevertheless, such a fibrous structure should significantly increase the surface area of the hybrid layer for electrochemical reactions and could be the explanation for the improved electrochemical response for the [Fe(CN)_6_]^3−^/[Fe(CN)_6_]^4−^ couple observed previously in this study.

**Fig. 4.**
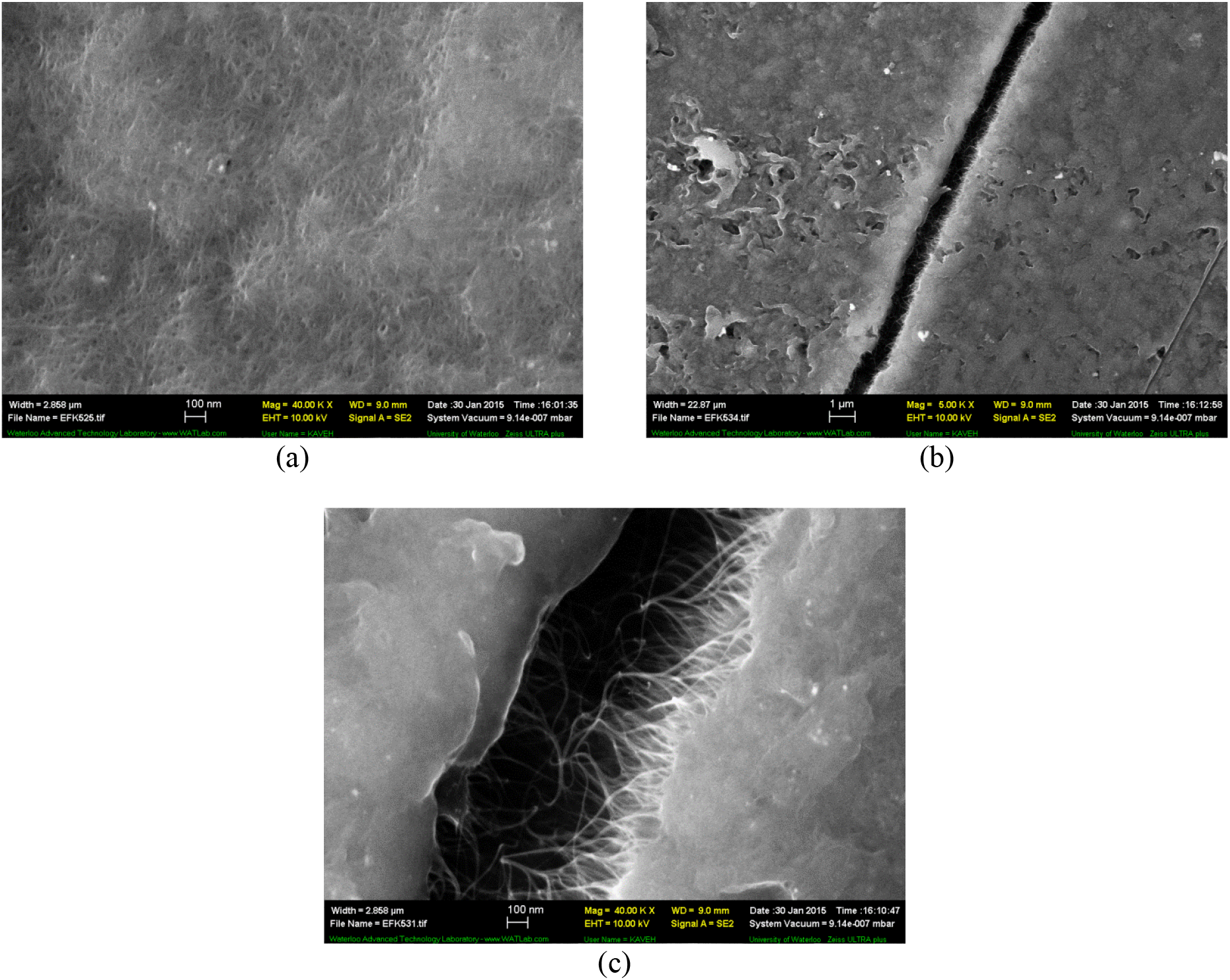
(a) Top view of EFK8-SWNT coating. (b) Top view of coating after removal from the GCE surface, showing the crack that had developed. (c) Higher magnification of (b) showing the fibrous structure of the coating.

### 3.2 Immobilization of hemoglobin

Given these promising results in using the EFK8-SWNT-modified GCE electrode, a solution containing 5 mg ml^−1^ Hb was mixed with EFK8-SWNT dispersion and applied to the GCE. Figure 5a shows an AFM image of the EFK8-SWNT coating with a fibrous structure on the GCE consistent with the SEM image in Figure 4a. Figure 5b reveals the presence of some globular particles (some of which are indicated by arrows) on the surface of the electrode modified with EFK8-SWNT-Hb. These particles which are not seen on the EFK8-SWNT-modified surface are likely bundles of Hb molecules given that each Hb molecule has a diameter of ~ 5 nm. To obtain further evidence that Hb had been immobilized on the surface, a CV obtained using the modified EFK8-SWNT-Hb-GCE electrode was compared to those obtained using electrodes that had not been contacted with Hb (i.e., EFK8-SWNT-modified GCE and bare GCE electrodes). In each case, the electrode was immersed in a 1M KCl solution containing 10mM K_3_Fe(CN)_6_. As shown in Figure 6a, the inclusion of Hb in the hybrid layer leads to a drop in the peak currents for the [Fe(CN)_6_]^3−^/[Fe(CN)_6_]^4−^ couple. Such an effect would be expected if Hb were successfully immobilized on the surface since this should lead to some inhibition of electron transfer. It is interesting to note that the current due to the [Fe(CN)_6_]^3−^/[Fe(CN)_6_]^4−^ couple rises more steeply and the peak separation is narrower on the EFK8-SWNT-Hb-GCE than on the bare GCE. This trend indicates faster electrode kinetics on the modified electrode than on the unmodified GCE although it remains considerably slower than on the EFK8-SWNT-GCE. EIS experiments were also conducted to check the consistency with the trends observed in the CVs. The data shown in Figure 6b reveal that the mixing of Hb with an EFK8-SWNT dispersion in preparing the coating for the GCE has a significant effect on the resulting Nyquist plots. The diameter of the high frequency semi-circle corresponding to the polarization resistance is larger than that obtained on the EFK8-SWNT-modified GCE. At the same time, it is still lower than that of the bare GCE which reflects the effect of SWNTs in facilitating electron transfer. The trends are perfectly in accord with those observed in the CVs in Figure 6a.

**Fig. 5.**
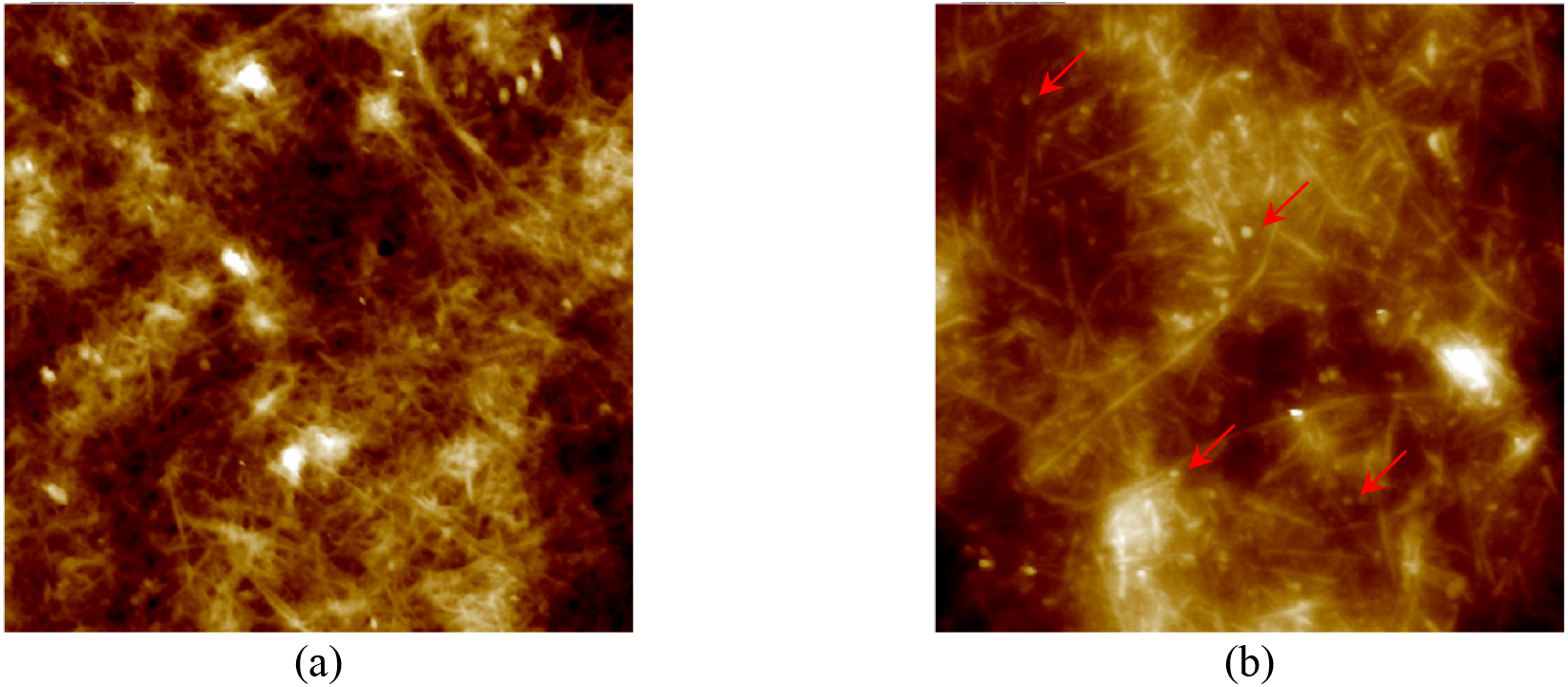
1μm×1μm AFM images of (a) EFK8-SWNT and (b) EFK8-SWNT-Hb coated GCE. Arrows in (b) show some globular Hb molecules.

**Fig. 6.**
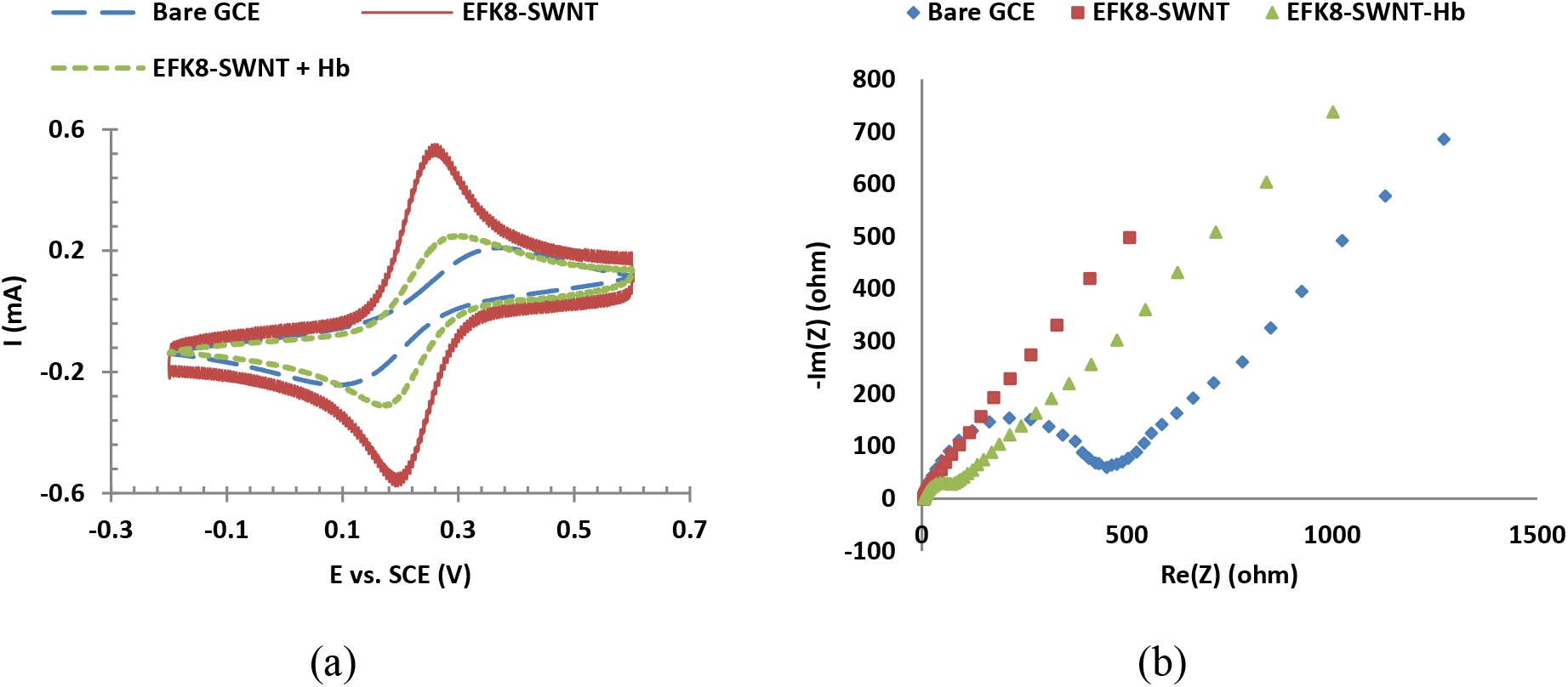
(a) CV and (b) Nyquist plots obtained on bare GCE, EFK8-SWNT and EFK8-SWNT-Hb-modified GCE immersed in 1M KCl containing 10mM K_3_Fe(CN)_6_.

### 3.3 Direct electrochemistry of hemoglobin on the electrode

To explore the electrochemical behavior of immobilized Hb on the surface, CVs were obtained on the modified GCEs in 0.1M PBS. Figure 7a shows that the presence of Hb on the modified GCE surface leads to higher cathodic current during the cathodic scan in which Fe^3+^ is reduced to Fe^2+^. This again supports the conclusion that Hb has not only been successfully immobilized on the surface but has also maintained electrochemical functionality, suggesting that it has not denatured and has retained its structure. In addition, it shows that the EFK8-SWNT modification layer provides a platform to utilize Hb in the direct electron transfer and mediatorless electron transfer on an electrode surface. As shown in Figure 7b, the current obtained on an EFK8-SWNT-Hb modified GCE is found to rise as the scan rate is increased from 20 to 200 mV s^−1^.

**Fig. 7.**
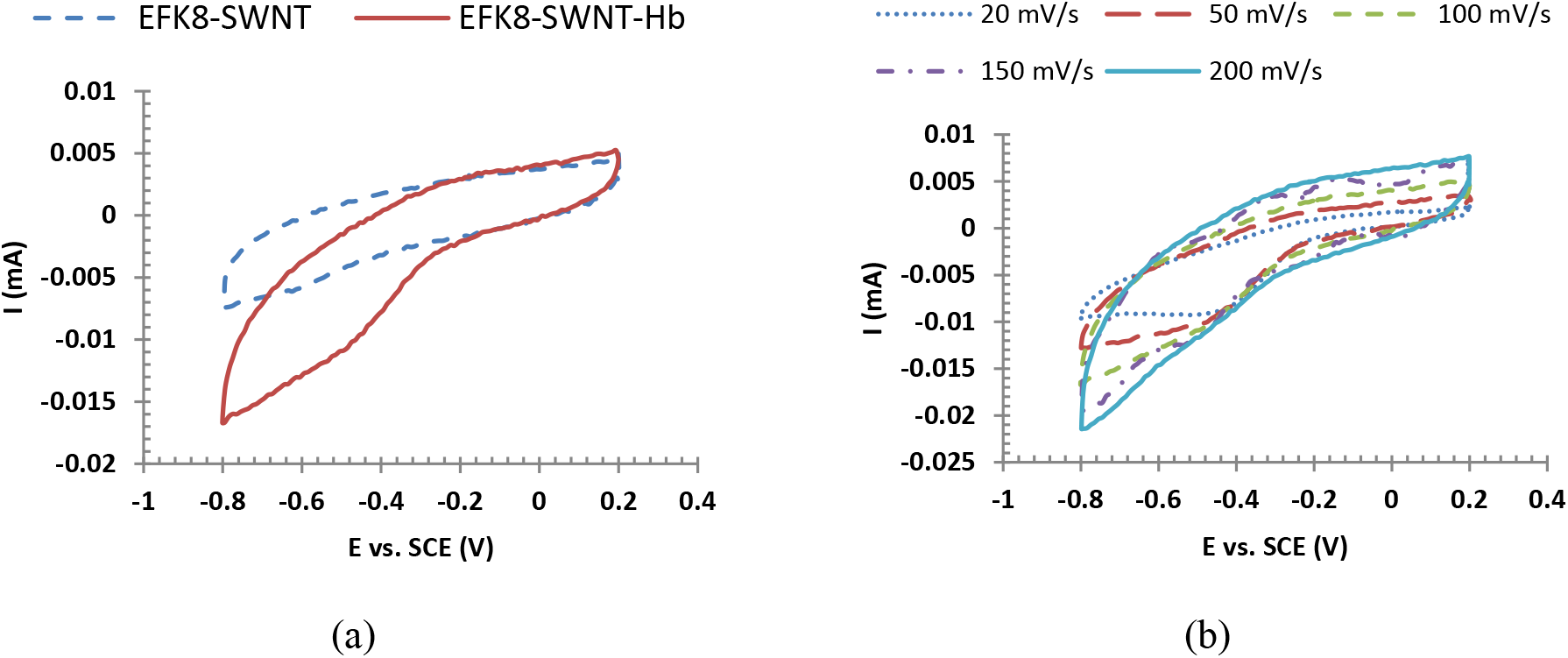
(a) CVs obtained on mediatorless EFK8-SWNT and EFK8-SWNT-Hb modified GCE in 0.1M PBS. (b) Effect of scan rate on CVs obtained on mediatorless EFK8-SWNT-Hb modified GCE in 0.1M PBS.

### 3.4 Biocatalytic activity of Hb within EFK8-SWNT

The bioelectrocatalytic activity of Hb for hydrogen peroxide reduction was investigated using CV and chronoamperometry. Only one EFK8-SWNT hybrid layer was applied to the GCE in this series of experiments. Figure 8 shows that the addition of 2.4 x 10^−4^ M H_2_O_2_ to PBS leads to a change in the voltammogram and the appearance of cathodic and anodic peaks. This shows that a GCE modified with EFK8-SWNT-Hb is sensitive to H_2_O_2_ and the immobilized Hb can catalyze reduction and oxidation of H_2_O_2_ without using any mediator. Furthermore, the current at a potential of −0.4 V (SCE) where H_2_O_2_ reduction occurs [29] increases as the H_2_O_2_ concentration rises. This sensitivity of the current to the H_2_O_2_ concentration makes EFK8-SWNT a promising platform to immobilize Hb and serve as a mediatorless biosensor to measure H_2_O_2_ concentrations.

**Fig. 8.**
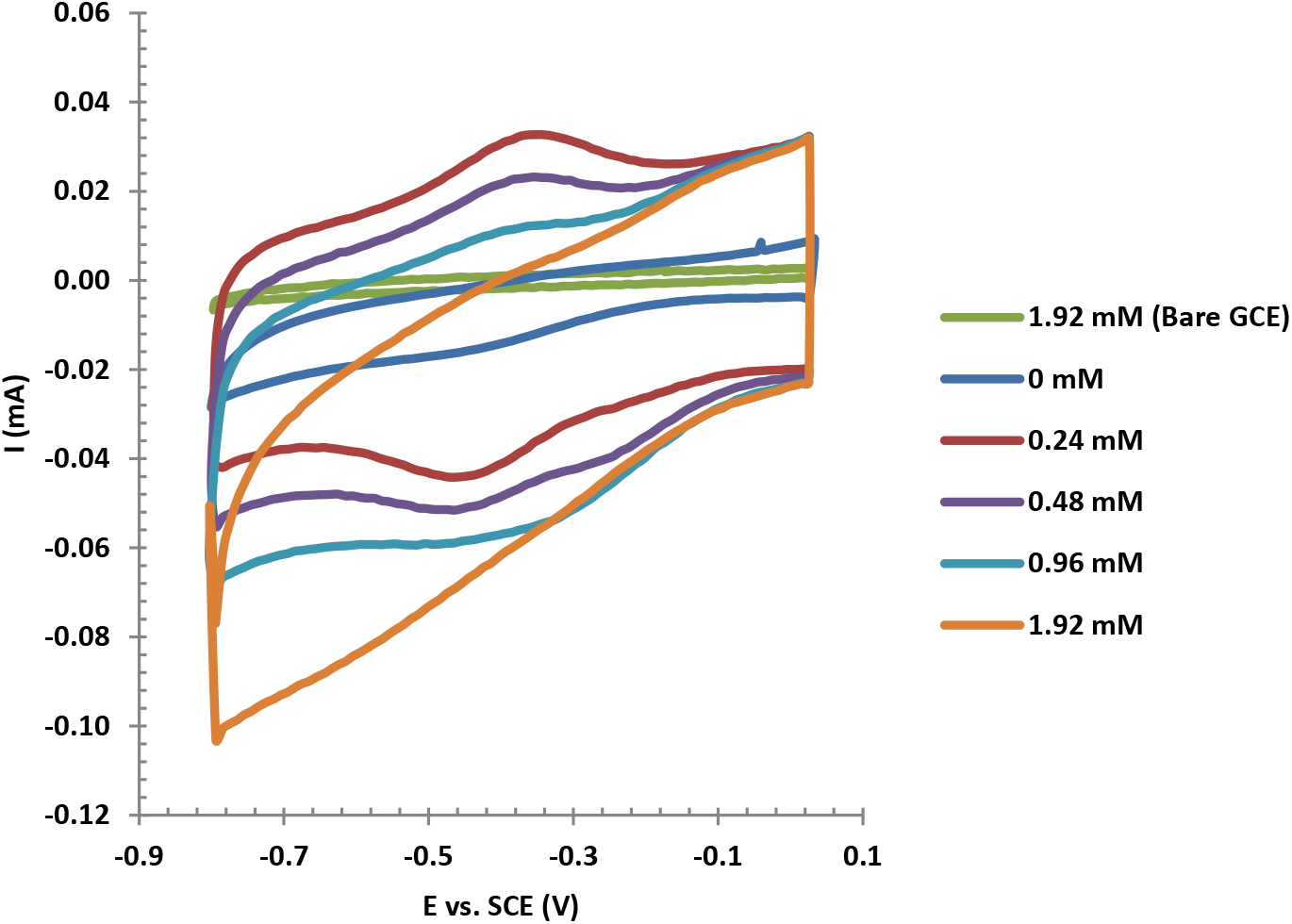
CVs of EFK8-SWNT-Hb-modified GCE in 0.1M PBS (pH 7.0) obtained at a scan rate of 100 mV s^−1^ and different H_2_O_2_ concentrations

To investigate this question more closely and quantitatively, a series of chronoamperometry experiments were conducted on an EFK8-SWNT-Hb-modified GCE to obtain a calibration curve of the current measured at −1.0V as a function of H_2_O_2_ concentration. Again, the GCEs were coated with only one EFK8-SWNT hybrid layer. In all cases, the current was found to reach steady state within 10 s. The calibration curve over the relatively wide range from 0 to 0.002 M H_2_O_2_ is presented in Figure 9a. The relationship between current and concentration is linear up to a concentration of 0.001 M before deviating above this level. Fig. 9b shows the calibration curve over the linear portion from 0 to 0.001 M. The high *r*^2^ value for this correlation and the relatively high sensitivity of 1.06 mA mM^−1^ cm^−2^ compared with that reported in previous work [2] shows that the EFK8-SWNT-Hb-modified GCE is promising for biosensor application. The calibration curve at very low concentrations between 0 and 0.0001 M is shown in the Figure 9c. Although the sensitivity of 0.639 mA mM^−1^ cm^−2^ is not as good as in Figure 9b, the calibration curve is still linear. Given that the cytotoxic level of H_2_O_2_ in the body is more than 50 μM [39], the results here demonstrate the effectiveness of this electrode as a H_2_O_2_ biosensor for clinical applications.

**Fig. 9.**
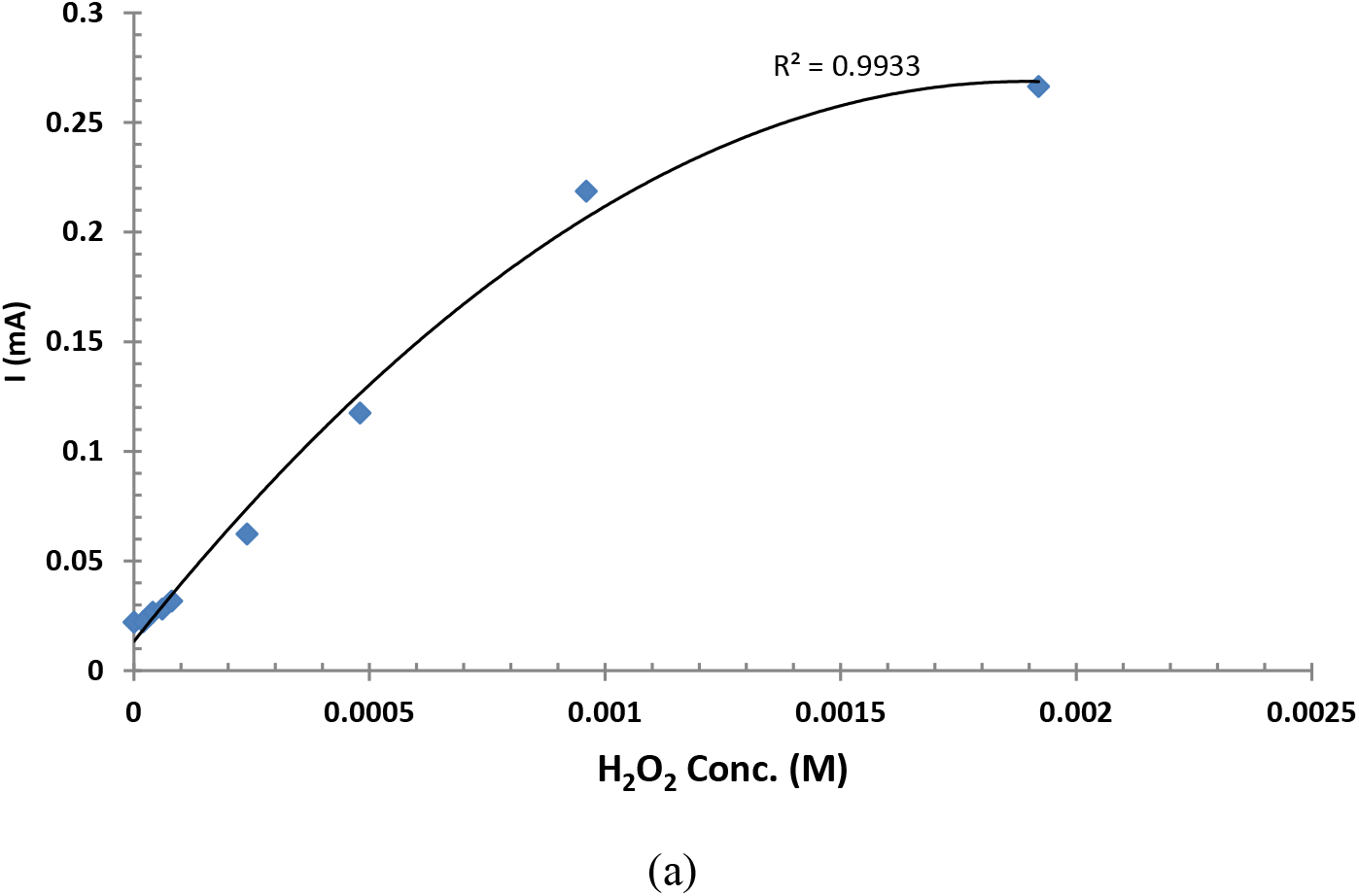
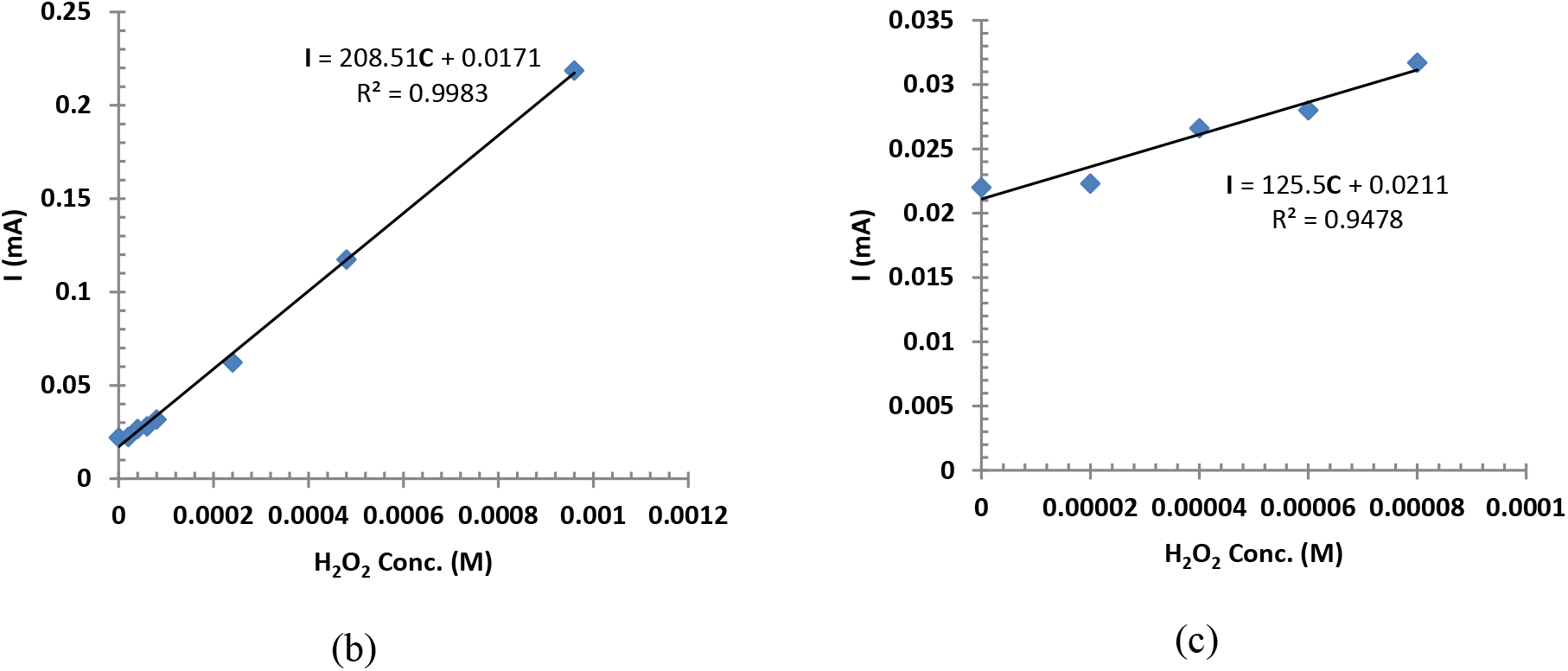
(a) Amperometric calibration curve of EFK8-SWNT-Hb-modified GCE at different H_2_O_2_ concentrations acquired at −1.0V after 10s. (b) Linear part of the graph. (c) Lower concentrations of the linear part.

Finally, the stability of the modified electrode was evaluated by measuring the current at a potential of −1.0V while being immersed in PBS (pH 7.0) at 4°C over a 14-day period. Although not shown here, the electrode retained 90.2% of its initial response to H_2_O_2_ during these amperometric measurements.

## 4. Conclusions

This study showed for the first time the use of dispersions of peptide and single-walled carbon nanotubes to modify GCEs to enhance their electrochemical response. It was shown that a peptide-only layer hinders the electrode response for the [Fe(CN)_6_]^3−^/[Fe(CN)_6_]^4−^ redox couple, whereas the inclusion of SWNTs in the same coating increases the electrode response significantly. This is attributed to the effect of the SWNTs in conducting electrons at the surface. We showed that hemoglobin could be easily immobilized on the electrode surface by mixing with the EFK8-SWNT dispersion and applying to the electrode. The immobilized Hb did not denature and was found to participate in direct electron transfer (presumably between the heme group and the electrode) in a PBS solution without the need for a mediator. This was again possible due to the role of SWNTs in tunneling electrons to the electrode surface. Furthermore, the modified electrode was shown to be sensitive to hydrogen peroxide dissolved in PBS and a promising mediatorless biosensor to measure its concentration. Based on these results, this modification provides a suitable platform for electrochemical mediatorless biosensors with redox proteins.

## Acknowledgments

The authors appreciate funding from the Natural Sciences and Engineering Research Council of Canada (NSERC) and the Canada Foundation for Innovation (CFI) to carry out this research. In addition, support from the Canada Research Chairs (CRC) program for one of the coauthors (P.C.) is gratefully acknowledged.

